# WISH-barcoding of *Salmonella* Typhimurium ATCC14028s strains for population dynamics studies *in vivo*

**DOI:** 10.64898/2026.04.29.721810

**Authors:** Christopher Schubert, Jeongmin Kim, Nicolas Näpflin, Melanie Hoos, Jemina Huuskonen, Christian von Mering, Wolf-Dietrich Hardt

## Abstract

**Background:** Barcoding of isogenic strains is a powerful approach to assess pathogen population dynamics during infection. Here, we adapted WISH-barcoding to *Salmonella* Typhimurium ATCC14028s to evaluate its suitability for pooled infection experiments in streptomycin-pretreated mouse models.

**Results:** WISH-barcoded wild-type pools showed pronounced population instability, characterized by stochastic strain loss and segregation into high- and low-fitness subpopulations. Whole-genome sequencing identified recurrent mutations in the methyltransferase *rsmG* and loss of the P3 plasmid carrying streptomycin resistance in low-fitness strains; neither was observed in Δ*invG* or Δ*ssaV* pools. We propose that *rsmG* mutations were enriched during strain construction carried out under streptomycin selection, following loss of the P3 plasmid. Control experiments demonstrated that *rsmG* mutations and P3 loss are counter-selected *in vivo* and attenuate gut-luminal colonization in streptomycin-pretreated mice.

**Conclusion:** While population dynamics experiments with ATCC14028s are feasible in principle, wild-type strains are prone to acquiring fitness-altering mutations during *in vitro* construction when using the P3 plasmid and streptomycin, highlighting the need for careful pool validation prior to use.

## Introduction

The gut microbiota confers colonization resistance against invading pathogens through various mechanisms, including nutrient competition (exploitation) and the secretion of antimicrobial compounds and Type VI secretion systems (interference) [1, 2]. Despite these barriers, *Salmonella enterica* serovar Typhimurium (*S*. Typhimurium), a Gram-negative enteric pathogen, has evolved strategies to circumvent these defenses, causing foodborne gastroenteritis in humans [3]. As a major cause of diarrheal infections worldwide, *S*. Typhimurium poses a significant public health concern [4]. To investigate the genetic requirements essential for *S*. Typhimurium colonization, genome-wide randomly barcoded transposon sequencing (RB-TnSeq) has previously been employed [5-8]. Transposon mutagenesis works by randomly inserting a transposon into the genome to disrupt genes, allowing identification of genes required for fitness or survival by tracking which insertions are lost under selective conditions [9]. This randomness can be a drawback when focusing on a specific physiological process, such as nutrient utilization, particularly when immune responses impose pronounced bottlenecks on the pathogen population [10, 11].

To address this limitation, the Wild-Type Isogenic Standardized Hybrid (WISH) barcode, a neutral genetic tag used to track individual strain abundance, was recently established [12] and adapted for *S*. Typhimurium SL1344 [11, 13-16]. Here, we aimed to adapt the WISH-barcoding system to ATCC14028, one of the two most widely used laboratory reference strains for *S*. Typhimurium virulence research. A key requirement is stable colonization, which is essential for robust assessment of mutant fitness in competitive index experiments. To quantify colonization stability, we calculate the Shannon Evenness Index (SEI), a metric of community evenness that captures how uniformly individual strains are represented within a biological sample, ranging from 0 (strong dominance by a single strain) to 1 (perfectly even distribution across strains) [17, 18]. A low SEI reflects the presence of population bottlenecks, such as the granulocyte-dependent bottleneck observed during *S*. Typhimurium infection in the streptomycin-pretreated mouse model between days 2 and 3 post-infection [10]. Similarly, in gnotobiotic mouse models, population bottlenecks occur alongside an increase in the inflammatory response, which is typically observed around day 4 post-infection [11]. Critically, beyond a certain threshold, genuine fitness differences become indistinguishable from stochastic effects imposed by population bottlenecks. Thus, SEI is a key parameter for assessing the robustness of fitness data.

For this reason, we used WISH-barcoded ATCC14028s pools to evaluate colonization stability in the streptomycin-pretreated mouse model. Experiments were performed in two specific pathogen-free (SPF) mouse strains, C57BL/6J and 129S6/SvEvTac. A key distinction between these models is the Nramp1 (Slc11a1) genotype: C57BL/6J mice carry a non-functional allele, rendering them highly susceptible to systemic *S*. Typhimurium infection, whereas 129S6/SvEvTac mice harbor a functional Nramp1 allele that confers increased resistance to systemic dissemination [19]. In addition to the wild-type strain, we analyzed *invG*- and *ssaV*-deficient ATCC14028s WISH-barcoded pools. InvG is a structural component of the Type III Secretion System 1 (T3SS-1) and is essential for host cell invasion [20, 21], whereas SsaV is a core component of T3SS-2, which is required for intracellular survival and systemic dissemination [22, 23]. By comparing Δ*invG* and Δ*ssaV* mutants with the wild-type pool, we assessed whether defects in epithelial invasion or systemic survival alter *S*. Typhimurium population stability in the streptomycin-pretreated mouse model during gut-luminal colonization.

## Results

### Construction of WISH-barcoded ATCC14028s pools to assess colonization stability

Throughout this study, we used the smooth strain ATCC14028s because it retains virulence in mice [24]. To assess the colonization stability of *S*. Typhimurium ATCC14028s in a streptomycin-pretreated mouse model, we rendered the parental strain streptomycin resistant by introducing the P3 plasmid from SL1344 via electroporation prior to WISH barcoding [25]. We then WISH-barcoded wild-type *S*. Typhimurium ATCC14028s with 10 unique WISH-tags. These tags were coupled to a carbenicillin/ampicillin resistance cassette, enabling quantification of the total tagged population by plating. Individual strain abundances in biological samples were determined by amplicon sequencing [11, 12]. In addition to the wild-type parental strain, we created 10 WISH-barcoded strains of Δ*invG* and Δ*ssaV* mutants in the ATCC14028s background (**Fig 1a**). We used two streptomycin-pretreated mouse models, specific pathogen free C57BL/6J and 129S6/SvEvTac mice. In both models, mice were treated with 25 mg streptomycin prior to infection to reduce the microbiota, allowing *S*. Typhimurium to reproducibly colonize the gut [20]. On the day of infection, mice were inoculated with 5 × 10^7^ CFU by orogastric gavage. Fecal samples were collected for three consecutive days post-infection. On day three, mice were euthanized and dissected, and in addition to fecal samples, cecal content and systemic organs - including the mesenteric lymph nodes, liver, and spleen - were collected (**Fig 1b**). To assess the colonization stability of ATCC14028s, all samples, except for organs, were enriched in selective LB (100 μg/ml carbenicillin) at 37°C for 4 hours. Afterwards, genomic DNA (gDNA) was isolated, and the library for amplicon sequencing was prepared through two consecutive PCR reactions (**Fig 1c**). WISH-barcodes were quantified by amplicon sequencing and analyzed using the mBARq software suite [26], and the resulting counts were used to calculate SEI values (**see Methods, Supplementary Data 1 and 2**).

**Figure 1.**
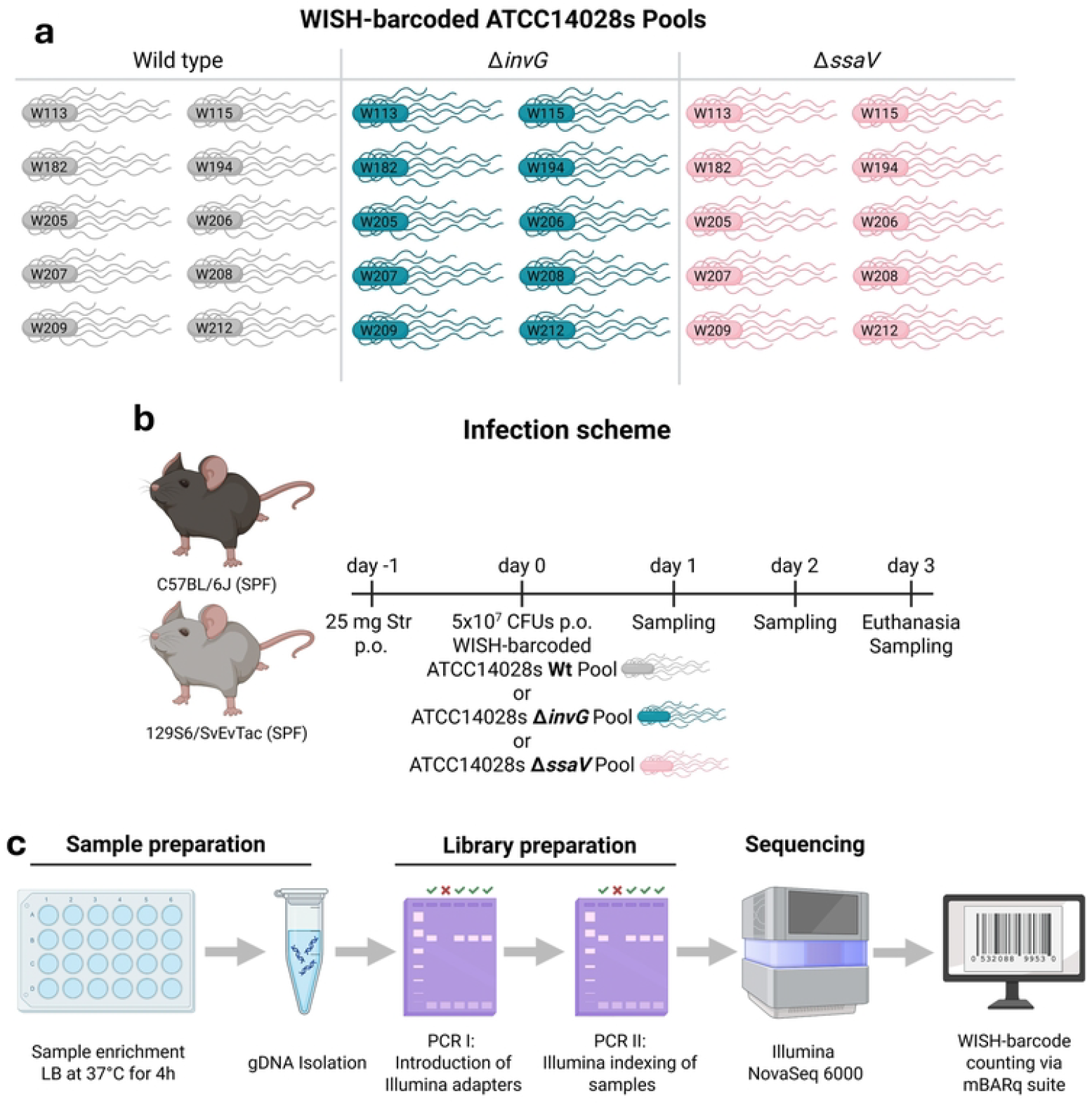
Experimental design of WISH-barcoded ATCC14028s pool infections. (**a**) Composition of WISH-barcoded ATCC14028s pools. The wild-type, Δ*invG*, and Δ*ssaV* pools each consisted of 10 uniquely WISH-barcoded strains. Created in BioRender. Schubert, C. (2026) https://BioRender.com/8twomu6 (**b**) Infection scheme. Streptomycin-pretreated C57BL/6J and 129S6/SvEvTac mice were orally inoculated with 5 × 10^7^ CFU of the indicated WISH-barcoded pools. Fecal samples were collected over three days post-infection, followed by euthanasia and sampling of feces, cecal content, and organs, including liver, spleen, and mesenteric lymph nodes. Created in BioRender. Schubert, C. (2026) https://BioRender.com/8twomu6 (**c**) Library preparation workflow. Samples were enriched in selective LB medium, genomic DNA was isolated, Illumina adapters and sample indices were introduced by PCR, and WISH barcodes were quantified by amplicon sequencing to determine relative strain abundances. Created in BioRender. Schubert, C. (2026) https://BioRender.com/8twomu6

### ATCC14028s wild-type populations show high degrees of unevenness when colonizing the gut of streptomycin-pretreated mice

To assess the evenness of gut-luminal ATCC14028s colonization, we compared WISH-barcoded wild-type ATCC14028s pools alongside Δ*invG* and Δ*ssaV* pools (**Fig. 1a**). Streptomycin-pretreated C57BL/6J and 129S6/SvEvTac mice were inoculated with 5 × 10^7^ CFU, and fecal samples were collected over three days post-infection. The evenness of the gut-luminal population was quantified using the Shannon Evenness Index (SEI). A conservative cutoff of 0.9 has been established previously, with samples exhibiting an SEI ≥ 0.9 considered stably colonized [10, 27]. In both mouse models, infection with the WISH-barcoded *S*. Typhimurium wild-type pool resulted in a significantly lower SEI compared with the Δ*invG* and Δ*ssaV* pools. In contrast, the SEI of the gut-luminal Δ*invG* and Δ*ssaV* pools remained stable throughout the three days of infection. The Δ*ssaV* pool consistently exhibited significantly higher SEI values in both mouse models than the Δ*invG* pool (**Fig. 2a, b**). Nevertheless, the SEI values of both Δ*invG* and Δ*ssaV* pools were sufficiently high to enable robust assessment of population dynamics using WISH-barcoding.

**Figure 2.**
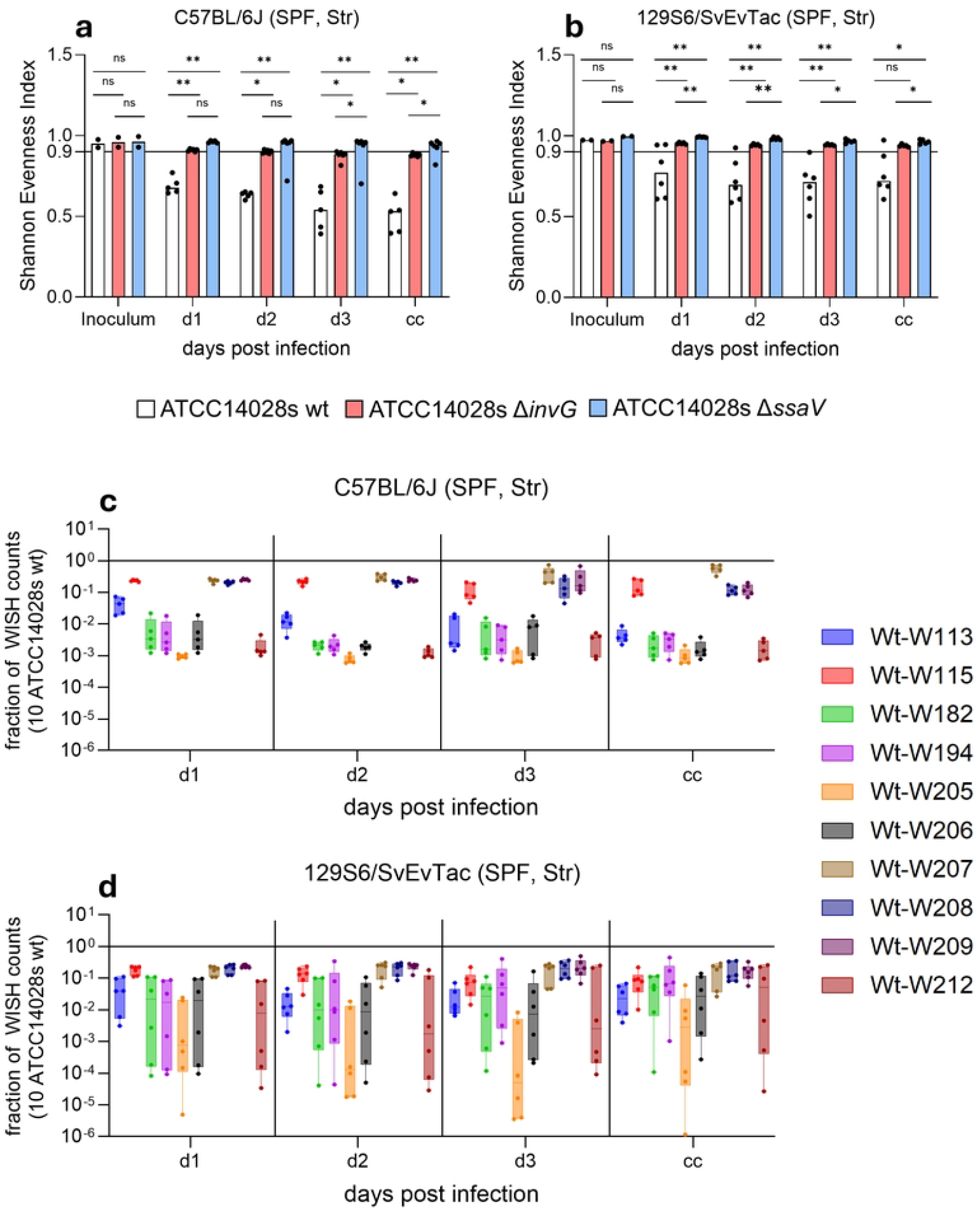
Δ*invG* and Δ*ssaV* mutants have a significantly higher SEI in both mouse models. The Shannon Evenness Index (SEI) was calculated for WISH-barcoded ATCC14028s strains across the three pools: wild type, Δ*invG*, and Δ*ssaV*. SEI values for the streptomycin-pretreated C57BL/6J (**a**) and 129S6/SvEvTac (**b**) models are presented as bar plots, displaying the median and individual data points. A threshold SEI value of 0.9 is indicated. Panels (**c, d**) depict the fraction of individual WISH counts normalized to the total WISH counts across all 10 WISH barcodes per pool. The respective mouse model is indicated above each diagram. Both panels are presented as box-and-whisker plots, showing the median, interquartile range (25th to 75th percentiles), minimum and maximum values, and individual data points. P values were determined using the two-tailed Mann–Whitney U-test: ** ≙ P < 0.005; * ≙ P < 0.05; ns ≙ P > 0.05.

To further assess why ATCC14028s wild-type pools yielded lower SEI values, we plotted the relative abundance of each WISH-barcode across all days post-infection, including cecal samples (**Fig. 2c, d**). In the C57BL/6J model, a distinct separation into two clusters was observed: Four WISH-barcodes with a high relative abundance and six with a lower relative abundance. Notably, all data points are clustered consistently, indicating negligible mouse-to-mouse variability (**Fig. 2c**). If a WISH-barcode had a high relative abundance in any individual mouse, it remained dominant throughout the infection period. Interestingly, WISH-barcodes with a high relative abundance in the C57BL/6J model also exhibited a high relative abundance in the 129S6/SvEvTac model, whereas low-abundance WISH-barcodes showed noticeably more pronounced mouse-to-mouse variability (**Fig. 2d**). Importantly, all inocula exhibited SEI values >0.95, with each WISH-barcoded strain contributing approximately 9.5% of the total population in both mouse models (**Supplementary Fig. 1a, b**). Both experiments were performed independently using the same pre-mixed WISH-barcoded ATCC14028s cryostocks (**see Methods**). This suggests that the competitive advantage of specific WISH-barcoded strains arose before the *in vivo* experiment. We reasoned that this was likely attributable to mutations acquired during the construction of the WISH-barcoded ATCC14028s wild-type strains.

### RsmG partially explains population dynamics in the WISH-barcoded ATCC14028s population

We observed two subpopulations – a low-fitness and a high-fitness group – within the WISH-barcoded wild-type pool (**Fig. 2c, d**), raising the question of whether this difference in colonization resulted from mutations acquired during strain construction *in vitro*. We therefore sequenced all three pools — wild type, Δ*invG*, and Δ*ssaV* — including the parental strains. Interestingly, all WISH-barcoded ATCC14028s strains with attenuated fitness were lacking the P3 plasmid (**Supplementary Fig. 2**). It should be noted that streptomycin resistance is not necessarily required to colonize the streptomycin-pretreated mouse model [28, 29]. However, we have repeatedly observed that streptomycin resistance confers a significant competitive advantage in co-infection experiments, particularly when one strain lacks resistance in our streptomycin-pretreated mouse models. For this reason, we compared ATCC14028s carrying P3-conferred streptomycin resistance against ATCC14028s lacking P3 in a competition experiment and observed a fitness attenuation in the susceptible strain (**Supplementary Fig. 3**). This attenuation was consistent with the fitness defect observed in the pooled experiment. To further investigate this attenuation, we focused on mutations acquired in strains that had undergone P3 plasmid loss. Compared with the parental wild-type strain, four of six WISH-barcoded wild-types that exhibited reduced abundance carried mutations in *rsmG*, which were absent from the WISH-barcoded Δ*invG* and Δ*ssaV* pools (**Supplementary Data 3-5**). *rsmG* encodes a 16S rRNA methyltransferase that fine-tunes ribosome function and translational fidelity [30]. In Wt-WISH182 and Wt-WISH194, we identified Gly75Val and Gly77Arg substitutions, which replace small, flexible L-glycine residues with bulky amino acids, likely disrupting protein folding and potentially impairing RsmG function (**Fig. 3a**). In contrast, Wt-WISH206 carried a >10-bp deletion in *rsmG*, resulting in a frameshift mutation, whereas Wt-WISH212 contained a mutation introducing an early stop codon. In all cases, these mutations are predicted to disrupt RsmG function.

**Figure 3.**
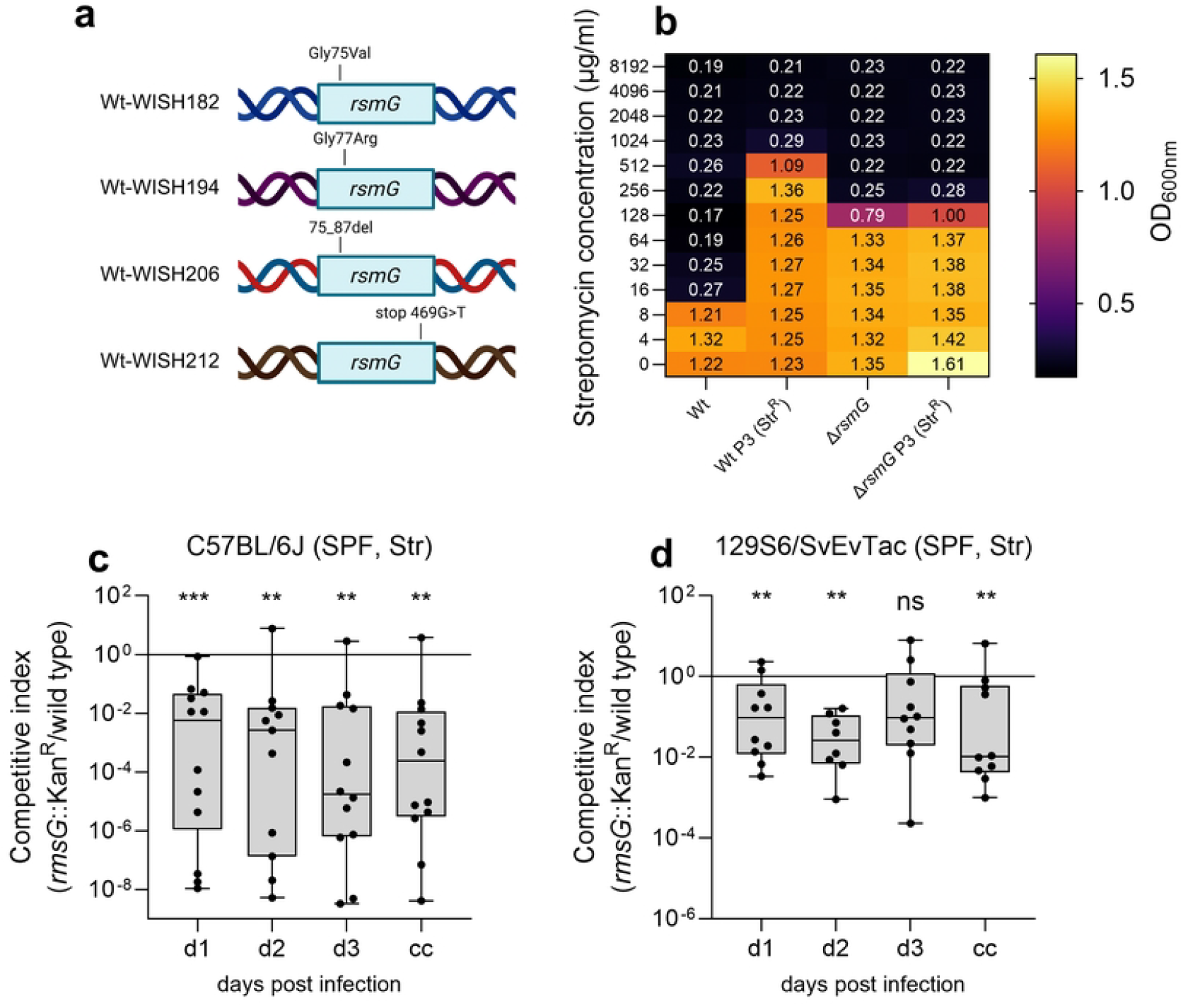
*rsmG* mutations have reduced fitness *in vivo*. (**a**) Schematic overview of mutations identified in *rsmG* in low-abundance WISH-barcoded wild-type strains, including missense substitutions (Gly75Val, Gly77Arg), a >10 bp deletion, and an early stop codon. Created in BioRender. Schubert, C. (2026) https://BioRender.com/8twomu6 (**b**) Minimal inhibitory concentration (MIC) assay for streptomycin showing growth (OD_600_) of wild type, wild type rendered streptomycin resistant with P3 plasmid (Wt Str^R^), *rsmG* mutant, and *rsmG* mutant with P3 plasmid across increasing streptomycin concentrations, indicating a modest effect of *rsmG* loss on streptomycin susceptibility. (**c, d**) Competitive indices of a *rsmG*::Kan^R^ mutant relative to wild-type ATCC14028s in streptomycin-pretreated C57BL/6J (n = 12) (**c**) and 129S6/SvEvTac (n = 10) (**d**) mice over the course of infection and in cecal content (cc). Each point represents one mouse; boxes indicate median and interquartile range. Statistical significance was assessed using the Wilcoxon rank-sum test (ns, not significant; **P < 0.01).

ATCC14028s is not naturally streptomycin resistant; however, *rsmG* mutations have been reported to reduce streptomycin susceptibility [30-32]. To test whether this phenotype applied here, we performed minimal inhibitory concentration (MIC) assays with ATCC14028s wild type and the *rsmG* mutant, as well as with both strains rendered streptomycin resistant by introduction of the P3 plasmid from SL1344 [25]. The MIC assay showed that the *rsmG* mutant exhibits a modest reduction in streptomycin susceptibility compared with the wild type, consistent with low-level tolerance rather than true resistance. The wild type exhibited an MIC of 16 μg/ml, compared with 256 μg/ml for the *rsmG* mutant. In contrast, introduction of the SL1344 P3 plasmid confers high-level streptomycin resistance only in the wild type (MIC of 1024 μg/ml Str). Interestingly, the introduction of the P3 plasmid did not alter the MIC of the *rsmG* mutant (**Fig. 3b**).

### Attenuation of the *rsmG* mutant in the streptomycin mouse model

Next, we sought to determine whether *rsmG* mutations could account for the fitness attenuation observed in ATCC14028s in our experiments. Rather than reconstructing each individual mutation, we assessed whether a *rsmG*-deficient strain exhibits a fitness defect during colonization. To this end, we generated an ATCC14028s *rsmG*::Kan^R^ mutant by P22 transduction and co-infected streptomycin-pretreated C57BL/6J and 129S6/SvEvTac mice with wild-type ATCC14028s and the *rsmG*::Kan^R^ mutant. In both mouse models, the *rsmG* mutant was consistently outcompeted by the wild type (**Fig. 3c, d**), in some cases to a degree comparable to that observed in the WISH-barcoded ATCC14028s wild-type pools (**Fig. 2c, d**). These results suggest that the *rsmG* mutations identified in our sequencing data may at least partially explain the reduced abundance of WISH-barcoded strains observed in the initial experiments (**Fig. 2**). Fitness effects of mutations are often condition dependent, such that a mutation conferring a growth advantage *in vitro* under streptomycin selection can impose a fitness cost during *in vivo* infection. This context-specific trade-off highlights how adaptations beneficial under artificial selection pressures may be detrimental within the complex environment of the mammalian host.

### Reduced fitness of the WISH-barcoded Δ*ssaV* pool at systemic sites

To assess whether the *invG* and *ssaV* mutations affected overall bacterial fitness, we quantified bacterial loads in feces, cecal content, and organs in streptomycin-pretreated C57BL/6J and 129S6/SvEvTac mice. Overall, bacterial loads in the feces and the cecum content were comparable across all parental strains in both models (**Supplementary Fig. 4a, b**). In contrast, systemic dissemination differed between strains. In streptomycin-pretreated C57BL/6J mice, the Δ*ssaV* pool exhibited significantly reduced bacterial loads in mesenteric lymph nodes, spleen, and liver compared with the wild-type and Δ*invG* pools, with most liver samples near or below the detection limit (**Supplementary Fig. 4c**). A similar trend was observed in the 129S6/SvEvTac model, although differences did not reach statistical significance (**Supplementary Fig. 4d**). To verify inflammatory responses in the gut, we measured the fecal lipocalin-2 (LCN2) levels. In the C57BL/6J model, LCN2 levels were significantly lower in the mice infected with the Δ*invG* and Δ*ssaV* pools compared with the wild type, while in the 129S6/SvEvTac model, only minor, transient differences were observed (**Supplementary Fig. 4e, f**). Nevertheless, all strains exceeded the established inflammatory threshold of 500 ng/g feces, previously defined as indicative of colitis [33]. Together, these data indicate that disruption of *invG* or *ssaV* does not substantially impair intestinal colonization, although the Δ*ssaV* pool shows reduced systemic dissemination. The enhanced gut-luminal colonization stability of the Δ*ssaV* and Δ*invG* pools relative to the wild-type pool is attributable to the retention of the P3 plasmid and wild-type *rsmG*, rather than to differences in virulence or inflammatory responses.

## Discussion

We assessed population dynamics of WISH-barcoded ATCC14028s wild-type, Δ*invG*, and Δ*ssaV* pools, each comprising 10 uniquely WISH-barcoded strains (**Fig. 1a**). In streptomycin-pretreated C57BL/6J and 129S6/SvEvTac mouse models, we observed stochastic loss of individual strains within the wild-type ATCC14028s pool (**Fig. 2a, b**). Further analysis revealed a separation into four WISH-barcoded wild-type strains with higher fitness and six with reduced fitness during mouse gut colonization (**Fig. 2c, d**). Whole-genome sequencing revealed that low-fitness WISH-barcoded wild-type strains had lost P3 plasmid-mediated streptomycin resistance and acquired mutations in *rsmG*, a gene encoding a 16S rRNA methyltransferase. While *rsmG* mutations partially compensated for P3 plasmid loss by reducing streptomycin susceptibility *in vitro*, they conferred a significant fitness disadvantage during *in vivo* colonization — a cost that, combined with P3 plasmid loss, likely accounts for the overall reduction in competitive fitness observed in these strains (**Fig. 3c**,**d and Supplementary Fig. 3**). In contrast, no such population stratification or fitness defects were observed in the Δ*invG* or Δ*ssaV* pools (**Fig. 2a and Supplementary Data 4 and 5**). Importantly, neither *rsmG* mutations nor P3 plasmid loss were detected in the Δ*invG* or Δ*ssaV* strain pools. Notably, all pools - wild-type, Δ*invG*, and Δ*ssaV* - were constructed using the same workflow and within the same timeframe, effectively excluding methodological artifacts. This selective pressure appeared to be specific to the wild-type background, though the underlying reasons for this remain unclear.

Streptomycin resistance in *Mycobacterium tuberculosis* is frequently not explained by the classical *rrs* or *rpsL* mutations, prompting the identification of *gidB* (synonymous to *rsmG*) as an additional resistance locus [30]. *gidB* encodes a conserved 16S rRNA m^7^G methyltransferase (modifying G527), and loss-of-function mutations confer low-level streptomycin resistance. Genes such as *rsmG* appear among isolates at elevated frequency not because of increased intrinsic mutation rates, but because they tolerate a broad range of loss-of-function mutations without severely compromising viability [30]. Streptomycin exposure, as encountered during *in vitro* strain construction, likely enriches pre-existing *rsmG* variants in ATCC14028s, especially in the context of P3 plasmid loss, thereby providing a mechanistic explanation for the repeated and independent emergence of *rsmG* mutations observed across strains. However, our data demonstrate that both *rsmG* mutations and P3 plasmid loss are counter-selected during infection, as they impose a measurable fitness cost during *in vivo* colonization (**Fig. 3c, d and Supplementary Fig. 3**). Intriguingly, despite sharing the same experimental workflow, this behavior was observed only in the wild-type background; why the Δ*invG* and Δ*ssaV* strains behaved differently remains an open question. Importantly, these mutations were not present in the parental strain used to generate the 10 uniquely WISH-barcoded derived strains included in the pools (**Supplementary Data 3-5**). Thus, the mutations were likely acquired during P3-transformation or WISH-barcoding and subsequent *in vitro* culturing, which was partially performed under streptomycin selection.

High-throughput mutant pool approaches enable efficient assessment of colonization fitness but critically depend on stable parental strain behavior [9]. The WISH-barcoding system was recently established [12] and adapted for use in *S*. Typhimurium SL1344 [11, 13, 14]. Random strain loss has also been reported for SL1344, where it coincides with inflammation-associated bottlenecks in streptomycin-pretreated C57BL/6J mice, typically beginning at day 2 post-infection, and during late-stage infection (days 3–4 post-infection) in the gnotobiotic OligoMM^12^ model [10, 11]. Unlike ATCC14028s, SL1344 did not exhibit a tendency to acquire *rsmG* mutations or lose P3 plasmid during *in vitro* manipulation. This may reflect the fact that P3 is the native plasmid of SL1344, potentially conferring greater plasmid stability in this background compared to ATCC14028s, in which P3 was introduced exogenously. The WISH-barcoded ATCC14028s Δ*invG* and Δ*ssaV* pools exhibited more stable colonization dynamics, reflecting the retention of P3 plasmid and the absence of fitness-altering mutations in these strains. The stratification of the wild-type ATCC14028s pool into low- and high-fitness subpopulations is attributable to the inadvertent loss of P3 plasmid and acquisition of *rsmG* mutations during strain construction, both of which imposed a fitness cost during *in vivo* colonization.

Collectively, our findings reveal that ATCC14028s can readily undergo *in vitro* mutational changes that were not observed in the Δ*invG* or Δ*ssaV* backgrounds, leading to disrupted population evenness in WISH-barcoded pool experiments *in vivo*. Further work is required to fully understand the basis of this behavior in the ATCC14028s wild-type strain and why it is not observed in the Δ*invG* and Δ*ssaV* backgrounds. Regardless, our findings highlight the importance of validating parental strain stability prior to pooled fitness experiments. This is particularly critical when applying WISH-barcoding approaches that rely on P3-encoded streptomycin resistance in ATCC14028s to assess fitness in streptomycin-pretreated mice.

## Acknowledgements

We would like to acknowledge and thank the staff at the ETH animal facilities EPIC and RCHCI; especially Manuela Graf, Katharina Holzinger, Dennis Mollenhauer, Samuel Boateng, Sven Nowok, and Dominik Bacovcin, and extend many thanks to members of the Hardt lab, as well as the NCCR Microbiomes, for their helpful comments and discussions.

## Author contributions

C.S. and W.-D.H. conceived and designed the experiments. C.S. and J.K. performed the *in vivo* and *in vitro* experiments. M.H. constructed the *rsmG* mutant. J.H. constructed the Δ*invG* parental strain. N.N. and C.v.M. performed SNP analyses of the sequenced strains. C.S. wrote the manuscript with contributions from all authors.

## Data availability

The amplicon sequencing data from the WISH-barcoded *S*. Typhimurium experiments generated in this study, as well as the whole-genome sequencing data, including genomes, are available in the European Nucleotide Archive (ENA) under accession number PRJEB85516. We adhered to the data reuse guidelines presented in [34].

## Code availability

The code used for WISH-barcode counting is available on GitHub (https://github.com/MicrobiologyETHZ/mbarq)[26].

## Methods

### Ethics statement

All animal experiments were performed in accordance with the guidelines of the Kantonales Veterinäramt Zürich under licenses ZH158/19, ZH108/22 and ZH109/2022, and the recommendations of the Federation of European Laboratory Animal Science Association (FELASA).

### Animals

We used male and female mice aged 8-12 weeks and animals of either sex were randomly assigned to the experimental groups. The consideration of animal sex was not included in the study design. All mice were maintained on the normal mouse chow (Kliba Nafag, 3537; autoclaved; per weight: 4.5% fat, 18.5% protein, 50% carbohydrates, 4.5% fiber). The mice originated from C57BL/6J or 129S6SvEv/Tac breeders initially obtained from Jackson Laboratories. Mice with a normal complex microbiota were specific pathogen-free (SPF) and bred under full barrier conditions in individually ventilated cage systems at the EPIC mouse facility of ETH Zurich (light/dark cycle 12:12 h, room temperature 21±1°C, humidity 50±10%). All studies were conducted in compliance with ethical and legal requirements and were reviewed and approved by the Kantonales Veterinäramt Zürich under licenses ZH158/19, ZH108/22 and ZH109/2022.

### Bacteria and culture conditions

All *Salmonella* strains used in this study were derived from the smooth variant *Salmonella* Typhimurium ATCC14028s [24].They are listed in **Supplementary Data 6**. All strains were routinely grown overnight at 37°C in Lysogeny broth (LB) with agitation. Strains were stored at -80°C in peptone glycerol broth (2% w/v peptone, 5% v/v glycerol (99.7%)). Custom oligonucleotides were synthesized by Microsynth and are listed in **Supplementary Data 7**; plasmids used in this study are listed in **Supplementary Data 8**.

### Homologous recombination by Lambda Red

Single-gene knockout strains were generated using the lambda-red single-step recombination protocol [35]. Primers were designed with ∼40 bp homology arms targeting the genomic region of interest and a 20 bp priming region corresponding to the antibiotic resistance cassette. PCR amplification was performed using plasmid pKD13 as the template for kanamycin resistance (**Supplementary Data 8**). DreamTaq Master Mix (Thermo Fisher Scientific) was used according to the manufacturer’s instructions. PCR products were purified using the NucleoSpin DNA purification kit (Macherey-Nagel). SL1344 harboring the pKD46 plasmid was cultured for 3 h at 30 °C to early exponential phase, followed by induction with L-arabinose (10 mM; Sigma-Aldrich) for 20 min. Cells were washed with ice-cold 10% (v/v) glycerol and concentrated 100-fold. Purified PCR products were introduced by electroporation (1.8 kV, 5 ms), followed by recovery in SOC medium (SOB premix; Roth GmbH, supplemented with 50 mM D-glucose) for 2 h at 37 °C, and plating on selective LB agar. Successful gene knockouts were confirmed by PCR and agarose gel electrophoresis, followed by Sanger sequencing (Microsynth AG). Antibiotic resistance cassettes were subsequently excised using FLP recombinase-mediated recombination [36].

### Homologous recombination by P22 phage transduction

The WISH-barcodes were introduced into ATCC14028s via P22 phage transduction from previously published SL1344 strains [11] (**Supplementary Data 6**). P22 phages carrying the respective WISH-barcodes were incubated with the recipient ATCC14028s strains (wild type, Δ*invG*, or Δ*ssaV* parental strains) for 15 min, followed by plating on selective LB agar plates. To ensure the removal of residual phages, transduced colonies were streaked twice consecutively on selective LB-agar plates overnight. To confirm the absence of P22 phage contamination, colonies were screened using Evans Blue Uranine (EBU) LB-agar plates (0.4% w/v glucose, 0.001% w/v Evans Blue, 0.002% w/v Uranine). WISH-tags were integrated at a fitness-neutral locus between the pseudogenes *malX* and *malY*, as previously described [37]. It is important to note that all parental ATCC14028s strains - wild type, Δ*invG*, and Δ*ssaV* - were rendered streptomycin resistant via transformation with the P3 plasmid prior to the introduction of the WISH-barcodes.

### Preparation of the ATCC14028s wild type, Δ*invG*, or Δ*ssaV* pools

Each strain from the WISH-barcoded ATCC14028s wild type, Δ*invG*, or Δ*ssaV* pools was inoculated in selective LB medium containing carbenicillin (100 μg/ml) and grown overnight at 37°C to maintain selection for the WISH-barcode. The overnight culture was then diluted 5% (v/v) into fresh LB medium without antibiotics and grown at 37°C for 4 h until reaching late exponential phase. At this stage, strains were pooled in equal volumes, centrifuged (4,500 × g, 4 °C, 15 min), and the resulting cell pellets were resuspended in peptone-glycerol medium to 10% (v/v) of the original volume and aliquoted into 100 μl cryotubes for storage at -80 °C. This approach enables rapid use of WISH-barcoded ATCC14028s pools, as 20 μl of the pooled stock can be subcultured in LB medium for only 4 h prior to infection.

### Animal experiments

For the ATCC14028s pool experiments, 8-to 12-week-old mice were inoculated with cells prepared from a 4 h subculture in LB medium sourced from the pooled cryo stocks. Bacteria were washed once with PBS, normalized to an optical density at 600 nm (OD_600_) of 2, and mice were infected by oral gavage with 5 × 10^7^ CFU in 50 μl. At the experimental endpoint (day 3 post-infection), animals were euthanized by CO_2_ asphyxiation. Fresh fecal pellets and whole cecal content were sampled in 500 μl PBS + metal bead using a TissueLyser (Qiagen). Samples were then briefly centrifuged (200 × g, room temperature, 2 min) to sediment large fecal debris, followed by dilution for assessment of bacterial population size on selective MacConkey agar plates. All dilutions were performed in 96-well microtiter plates.

### Sample preparation for amplicon sequencing and WISH-barcode quantification

After homogenization, fecal and cecal samples (125 μl) were enriched in 1 ml selective LB medium containing 100 μg ml^−1^ carbenicillin for 4 h at 37 °C to select for WISH-barcoded strains. Bacterial cells were then pelleted, the supernatant was discarded, and the pellets were stored at −20 °C. DNA was extracted from thawed pellets using a commercial kit (Qiagen DNeasy Mini Kit) according to the manufacturer’s instructions. For PCR amplification of WISH-barcodes, 2 μl of genomic DNA and 0.5 μM of each primer (WISH_Illumina_fwd and WISH_Illumina_rev; see **Supplementary Data 7**) were used with DreamTaq Master Mix (Thermo Fisher Scientific). The reaction was conducted with the following cycling program: initial denaturation step at (1) 95°C for 3 min followed by (2) 95°C for 30 sec, (3) 55°C for 30 sec, (4) 72°C for 20 sec for (5) 25 cycles, and a terminal extension step at (6) 72°C for 10 min. PCR products were column-purified using the NucleoSpin Gel and PCR Clean-up kit (Macherey-Nagel) according to the manufacturer’s instructions. We indexed the PCR products for Illumina sequencing by performing a second PCR with nested unique dual index primers using the following program: (1) 95°C for 3 min, (2) 95°C for 30 s, (3) 55°C for 30 s, (4) 72°C for 20 sec, (5) repeat steps 2-4 for 10 cycles, (6) 72°C for 3 min. Afterward, we assessed the indexed PCR product using gel electrophoresis (1% w/v agarose, 1xTAE buffer), pooled the indexed samples according to band intensity, and subsequently purified the library via AMPure bead cleanup (Beckman Coulter) before proceeding to amplicon sequencing. Amplicon sequencing was performed by BMKGENE (Münster, Germany). Samples were sequenced by BMKGENE (Münster, Germany) at 1 Gb output on the Illumina NovaSeq platform using 150 bp paired-end reads. Subsequently, the reads were demultiplexed and grouped by WISH-tags using mBARq software suite [26]. Misreads or mutations of up to five bases were assigned to the closest correct WISH-tag sequence. The WISH barcode counts for each mouse in every experiment are available in **Supplementary Data 1 and 2**. These counts were used to calculate the competitive fitness and Shannon Evenness Index (SEI).

### Competitive experiment

Streptomycin-pretreated C57BL/6J and 129S6/SvEvTac mice were infected with a 1:1 mixture of ATCC14028s wild type and the *rsmG* mutant, totalling 5 × 10^7^ CFU. The inoculum load of each strain was enumerated on selective MacConkey agar plates and used to normalize bacterial loads measured from day 1 to day 3 post-infection. The normalized bacterial loads were then used to calculate the competitive index (mutant divided by wild type). Statistical significance was assessed by comparing the absolute bacterial loads of the wild type and the *rsmG* mutant.

### Lipocalin-2 analysis of feces samples

Lipocalin-2 levels were measured in fecal samples homogenized in 500 μl PBS using an ELISA assay (DuoSet Lipocalin-2 ELISA kit, DY1857; R&D Systems) according to the manufacturer’s instructions. Fecal pellets were diluted 1:20, 1:400, or left undiluted, and concentrations were determined using Four-Parametric Logistic Regression curve fitting.

### Minimum inhibitory concentration (MIC) determination

MICs were determined by broth microdilution in 96-well microtiter plates. Antibiotics were prepared as sterile stock solutions and serially two-fold diluted in Mueller-Hinton broth 2 to generate the indicated concentration range. Overnight bacterial cultures were diluted into fresh medium and grown to mid-log phase, then adjusted to an inoculum of ∼5 × 10^5^ CFU ml^−1^ in the final assay volume. Wells were inoculated with bacteria and included a growth control (no antibiotic) and a sterility control (medium only). Plates were incubated statically at 37 °C for 16–20 h. MIC was defined as the lowest antibiotic concentration that prevented visible growth (no turbidity) compared with the growth control. Where indicated, growth inhibition was additionally quantified by measuring OD_600_ using a plate reader. MIC assays were performed with at least two independent biological replicates, and representative median MICs are reported.

### Single-nucleotide polymorphism (SNP) analysis of ATCC14028s pools

WISH-barcoded ATCC14028s wild-type, Δ*invG*, and Δ*ssaV* strains were grown independently overnight in selective LB medium, followed by genomic DNA isolation using a commercial kit (Qiagen DNeasy Mini Kit) according to the manufacturer’s instructions. Genomic DNA samples were submitted to BMKGENE (Münster) for whole-genome sequencing. Quality controlled reads were used to assemble genomes with Shovill v1.1.0 (Seemann T (2016) https://github.com/tseemann/shovill) and SKESA v2.5.1 [38] as implemented in Bactopia v3.2.0 [39] using default settings. SNP profiles (and Insertion/Deletions) were generated by mapping quality control reads against a *Salmonella enterica* reference genome. To this end, a reference genome for *Salmonella enterica* subsp. *enterica* serovar Typhimurium str. 14028 (accession CP001363.1, [24]), plasmids P1 (accession HE654724.1), P2 (accession NC_017718.1), and P3 plasmid (accession HE654726.1) was downloaded from NCBI (accessed: 2025.04.15). Snippy v4.6.0 (Seemann T (2015) https://github.com/tseemann/snippy) as implemented in Bactopia was then used to obtain SNP profiles for each strain using default settings against the reference genome and plasmids. Functional annotations of sequences, in which SNPs or insertions/ deletions were detected, were obtained with NCBI Prokaryotic Genome Annotation Pipeline v2025-05-06.build7983 [40]. SNP profiles were aggregated for all strains using Python 3.10.8. Sequencing depths and coverages were calculated with samtools v1.21 [41] and visualized with matplotlib v3.9.1 [42] and seaborn v0.12.2 [43].

### Statistical analysis

No statistical methods were used to predetermine sample sizes. For all mouse experiments, sample sizes of 5 or greater were used, consistent with those reported in previous publications [44]. Data collection and analysis were not performed blind to the conditions of the experiments. Where applicable, the two-tailed Mann-Whitney U test was employed to assess statistical significance, as specified in the figure legends. Statistical analyses were performed using GraphPad Prism 10 for Windows. *P* values were grouped as follows: ** **≙** *P* < 0.005; * **≙** *P* < 0.05; ns **≙** *P* > 0.05.

